# Inferring perturbation profiles of cancer samples

**DOI:** 10.1101/2020.12.10.419077

**Authors:** Martin Pirkl, Niko Beerenwinkel

## Abstract

**Motivation:** Cancer is one of the most prevalent diseases in the world. Tumors arise due to important genes changing their activity, e.g., when inhibited or over-expressed. But these gene perturbations are difficult to observe directly. Molecular profiles of tumors can provide indirect evidence of gene perturbations. However, inferring perturbation profiles from molecular alterations is challenging due to error-prone molecular measurements and incomplete coverage of all possible molecular causes of gene perturbations.

**Results:** We have developed a novel mathematical method to analyze cancer driver genes and their patient-specific perturbation profiles. We combine genetic aberrations with gene expression data in a causal network derived across patients to infer unobserved perturbations. We show that our method can predict perturbations in simulations, CRISPR perturbation screens, and breast cancer samples from The Cancer Genome Atlas.

**Availability:** The method is available as the R-package nempi at https://github.com/cbg-ethz/nempi.

## 1 Introduction

Cancer progression is often linked to alterations in driver genes (Bailey *et al.*, 2018). A mutation in a driver gene increases the probability of cancer development. These genes are often not functioning normally in cancer cells, but are inhibited or over-expressed. We call genes with this abnormal behaviour perturbed. Perturbations are hard to observe directly. However, some observable alterations such as mutations provide evidence for a gene perturbation. If the gene has a non-silent mutation, it is probable that its behaviour is perturbed. The combination of different molecular profiles is useful to identify perturbed genes. In general, however, not all different types of measurements are available. For example, if only gene expression data is available, the identification of perturbed genes due to mutations may be difficult. Even if the mutation profiles are available, they may not reveal all perturbed genes correctly due to measurement error causing false positive and false negative mutation calls. In the case of a true negative mutation call, the gene could still be perturbed in a different way, e.g., by micro RNA activity (Shivdasani, 2006; O’Brien *et al.*, 2018).

Identification of driver genes is important to characterize cancer types and help establish useful therapies. Especially knowledge about the genomic landscape can be helpful to establish successful treatments (Al-Lazikani *et al.*, 2012). Several methods deal with driver gene identification on a global scale. Some methods rely mainly on mutation data to derive driver genes for specific cancers (Lawrence *et al.*, 2013). Other methods include descriptive features of the genes (Tokheim *et al.*, 2016) or combine different data types (Hou and Ma, 2014; Dimitrakopoulos *et al.*, 2018; Hou *et al.*, 2018) available from, for example, The Cancer Genome Atlas (TCGA, http://cancergenome.nih.gov/, Network *et al.*, 2008). However, not only the identification of driver genes is important, but also which gene is perturbed in which cancer sample, especially when it comes to supporting treatment decisions based on these information. A gene can be a driver for breast cancer, but is only mutated in a few samples. Maybe it is also perturbed in other samples, but not mutated. It is also useful to know, which other genes are perturbed and in what combinations. Hence, we want to know the perturbation profiles of each tumor.

Inferring perturbation profiles can be viewed as a classification problem for each gene. A sample would either be classified as “gene X is perturbed” or “gene X is not perturbed”. For example, one can learn a classifier for each gene, which is possibly perturbed, based on gene expression profiles. Hence, this problem can be solved with supervised learning methods, such as, for example, support vector machines (Cortes and Vapnik, 1995; Honghai *et al.*, 2005; Yang *et al.*, 2012), neural networks (Nelwamondo *et al.*, 2007, Smieja *et al.*, 2018) or random forests (Pantanowitz and Marwala, 2009). Alternatively, data imputation methods can also be used to infer incomplete perturbation profiles (Shah, 2018; Azur *et al.*, 2011; Stekhoven and Bühlmann, 2011).

We developed a novel method called nested effects model-based perturbation inference (NEM*π*), which uses supervised learning to infer unobserved perturbations. We use a network approach based on gene expression data with samples labeled by their perturbations. We use the inferred network to learn the complete perturbation profiles of all samples. We iteratively optimize the perturbation profile and relearn the network until a convergence criterion is reached (Fig. 1).

**Figure 1:**
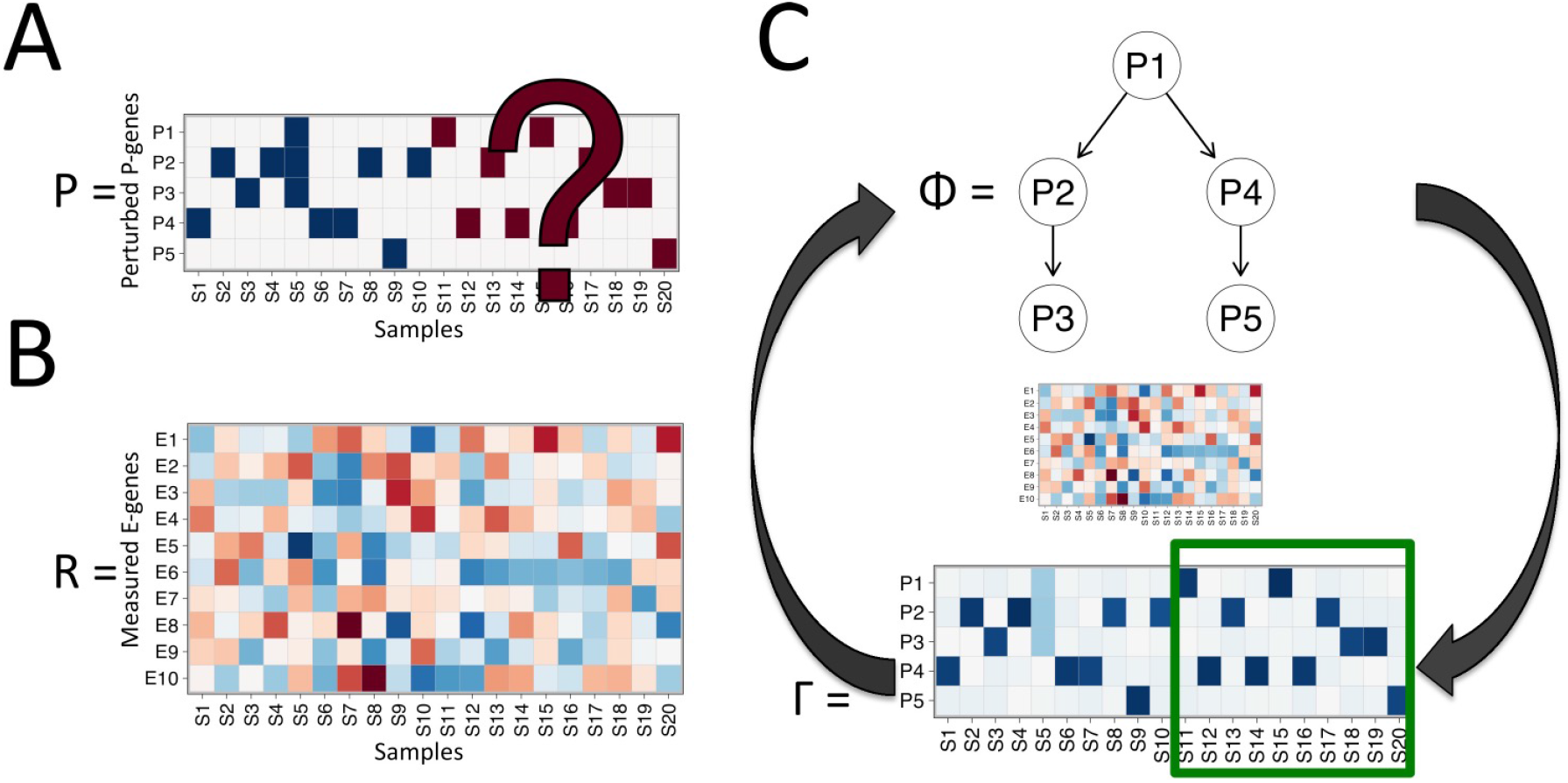
Perturbation inference scheme. The binary perturbation matrix *P* (A) with known (blue) and unknown (red) perturbations and the continuous log odds matrix *R* (B) derived from gene expression data *D* are available for the same set of samples. *P* and *R* are initially used to infer a causal network *φ* of the perturbed genes. Iterative EM algorithm (C): Based on *φ* and *R* the soft perturbation profile Γ is inferred. Γ and *φ* are iteratively updated until convergence. The incomplete part is inferred (green box) and the rest revised.

We validate NEM*π* on five single-cell RNA-seq (scRNA-seq) perturbation data sets from experiments using Clustered Regularly Interspaced Short Palindromic Repeats (CRISPR). These experiments are tailor-made to show that NEM*π* can successfully predict perturbations from gene expression data. Additionally, we perform an exploratory analysis on breast cancer (BRCA) data from TCGA. We use known mutation profiles to learn the perturbation profile of the patient samples. We compare the predicted perturbation profiles to copy number variations and methylated states of the corresponding mutated genes in the same patient samples.

## 2 Methods

NEM*π* is build on the causal network learning approach called Nested Effects Models (NEM, Markowetz *et al.*, 2005; Tresch and Markowetz, 2008). We extend this model to infer perturbation profiles from gene expression profiles.

NEM and its extensions have been applied to various perturbation data sets. Most recent versions of the algorithm have been extended to combinatorial perturbations (Pirkl *et al.*, 2016, Pirkl *et al.*, 2017) probabilistic perturbations (Srivatsa *et al.*, 2018), time-series data (Anchang *et al.*, 2009, Froehlich *et al.*, 2011, Wang *et al.*, 2014), hidden player inference (Sadeh *et al.*, 2013), context specific signaling (Sverchkov *et al.*, 2018) and single cell perturbations (Anchang *et al.*, 2018; Pirkl and Beerenwinkel, 2018). NEM*π* is related to NEMiX (Siebourg-Polster *et al.*, 2015). NEMiX also infers a perturbation. However, NEMiX predicts whether the whole pathway has been activated or not. Hence, it performs a clustering of samples (cells) into two clusters to account for inactive pathways explaining different expression profiles for the same perturbation of different samples. Unlike NEM*π*, NEMiX does not infer gene perturbations, but treats them as a prior fixed parameter.

### 2.1 Network model

Let *n* be the number of perturbed genes (P-genes) with unknown perturbation states in a subset of *u* samples. *m* is the number of features or effect genes (E-genes) for which gene expression data is available. Let *P* = (*p*_*ij*_) be the perturbation matrix with *p*_*ij*_ = 1, if P-gene *i* is perturbed in sample *j*. We assume that *P* is only known for a subset of samples.

We parameterize the causal network of the *n* P-genes by the transitively closed adjacency matrix *φ* of the P-genes. *θ* describes the relationship between P-genes and E-genes with *θ*_*ij*_ = 1, if P-gene *i* is the parent of E-gene *j*. We employ the assumption that each E-gene can have at most one parent. Our expected data pattern is computed by *F* = *φθ*. That is, if P-gene *i* is perturbed all descendants of *i* are also perturbed as well as all E-genes, which are children of the perturbed P-genes. Hence, *f*_*ij*_ = 1, if E-gene *j* is a child of *i* or a child of a descendant of *i*, and *f*_*ij*_ = 0 otherwise.

Let *D* = (*d*_*ij*_) be the gene expression data and *R* = (*r*_*ij*_) the corresponding log odds with

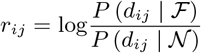

with the null model 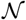, which does not predict any effects and the full model 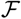, which predicts effects for all E-genes in all samples. I.e. 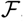 is the expected profile *F* of *φ*, if all P-genes are indistinguishable a nd a perturbation of one single P-gene causes a perturbation of all other P-genes. *r*_*ij*_ < 0 means E-gene *i* shows no effect in sample *j* and *r*_*ij*_ > 0 means E-gene *i* shows an effect in sample *j*. Hence large values in *R* correspond to the ones in *F*. As in Tresch and Markowetz (2008) we compute

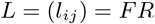

with *l*_*ij*_ the log odds of the perturbation of P-gene *i* in sample *j*. The full log odds for a candidate model (*φ, θ*) given the data can then be computed as

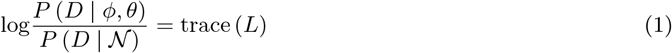

and is optimized with respect to *φ* and *θ*.

### 2.2 Perturbation inference

If the perturbation information is complete, we learn the causal network *φ* and E-gene attachments *θ* by optimizing the log odds in (1). However, in that case we assume *R* has the same number of columns (samples) than P-genes. In other words, for each P-gene we have a corresponding column in *R*, in which the P-gene is perturbed. In our more general case, we have many more samples than P-genes and allow for combinatorial perturbations. Because the perturbations are only observed in a subset of samples, we introduce the hidden random variable *Z* = (*z*_*ij*_) with *z*_*ij*_ = 1, if P-gene *i* has been directly perturbed in sample *j* and *z*_*ij*_ = 0 otherwise. The causal network *φ* propagates the direct perturbation to the descendants of P-gene *i*. This propagation is computed by Ω = *φ^T^ Z*. We call positive entries in *Z* direct perturbations, while Ω describes the actual perturbation profiles of the samples. For example, in Fig. 2 only P-gene 2 is directly perturbed in sample 7 (*z*_2,7_ = 1) and is also an ancestor of P-gene 3 (*φ*_2,3_ = 1). Hence, both P-genes 2 and 3 are perturbed in sample 7. Furthermore, each P-gene *i* has a prior probability *πi* = *P* (*z*_*ij*_ = 1) ∀*j* of being perturbed with

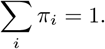

**Figure 2:**
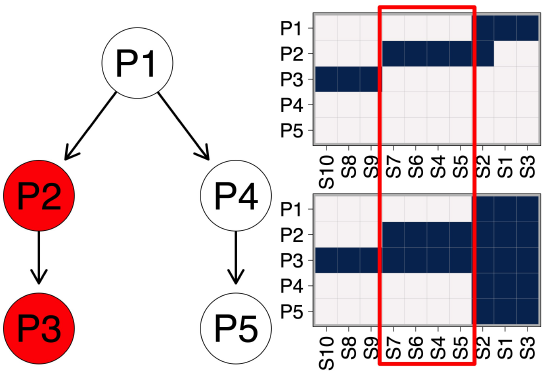
The network *φ* (left) predicts a perturbation of gene 3, if gene 2 is perturbed. Hence, the direct perturbation *Z* (top) does not need to include a perturbation of gene 3, since this is propagated via the network and included in the perturbation profile Ω = *φ^T^ Z* (bottom).

In our model the direct perturbations are not only propagated via the causal network *φ* but also via the E-gene attachments *θ*. Similar to before, this is done by matrix multiplication 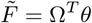. *Z* can have multiple 1s in each column and therefore we have to set values in 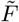, which are greater than 1 to 1. Hence, 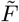, as previously *F*, describes the expected data pattern for all E-genes and samples in the log odds matrix *R*.

For maximum likelihood estimation, we want to know how probable are the gene expression profiles *D* given perturbation profiles *Z*. We need to maximize the probability of the full data (*D, Z*) given the model parameters, i.e., the causal network *φ* and the E-gene attachments *θ*,

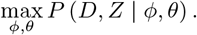

We re-formulate this optimization problem to maximizing the log odds

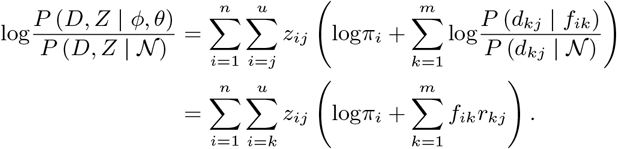

However, parts of the data are hidden (*Z*). We solve this problem by implementing an expectation maximization algorithm (Dempster *et al.*, 1977). In the E-step, we fix the causal network *φ*, the E-gene attachments *θ* and the P-gene priors *π* and compute the expectations of the direct perturbations *z*_*ij*_ for the *j*th sample *d*_*j*_ by

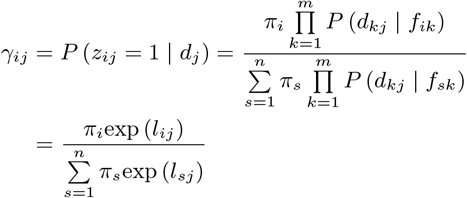

with Γ = (*γ*_*ij*_). In the M-step, we optimize the expected value of the log odds

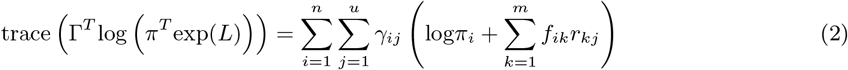

with respect to the causal network *φ* and the E-gene attachments *θ*. The priors *π* are computed by

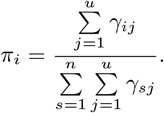

We perform the E- and M-step iteratively until the log odds in (2) or the parameters do not change anymore.

The optimization in the M-step is done by adding or removing edges in the causal network *φ*. All modifications of the network *φ* are evaluated before the change (greedy search). This is done until no change in *φ* increases the log odds anymore. After each change we are estimating the E-gene attachments *θ*. In each greedy search we can start with a specific network *φ*. More different starts increase the chances to reach a global optimum, but also increase the run time. We increase the probability that the log odds increase without using too much restarts by starting the greedy search three different times, from the previous solution *φ*_*i*−1_, the empty, and the fully connected network in the *i*th M-step with *φ*_0_ as the empty network. We take the highest scoring solution as the new causal network *φ*_*i*_. The run time complexity is 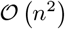 for every iteration in the number of P-genes *n*.

During the optimization of the M-step, we would also have to search for an optimal *θ*. However, we estimate the E-gene attachments *θ* in the following way. After changing an edge in *φ* and before computing the log odds, we estimate *θ* by first computing *Q* = *φ^T^* Γ*R^T^*, with *q*_*ij*_ as the log odds of the observed data pattern of E-gene *j* given that E-gene *j* is attached to P-gene *i*. We estimate the attachments *θ* by maximum a posteriori of the log odds *Q* with

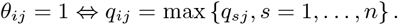

The priors *π* remain fixed during the optimization of the network *φ* and the E-gene attachments *θ*. Additionally we include a null E-gene, which does not predict any effect and has otherwise badly fitting E-genes attached to it.

To avoid over-fitting, we a dd a null component (null P-gene), which does not predict any effects. Hence, if the null gene dominates the log odds for a sample, it is hardly used in the inference. In other words, the model predicts that no P-gene was perturbed in the sample.

The initial expectation Γ_0_ of the direct perturbations *Z* is based on a given incomplete perturbation matrix *P* = (*p*_*ij*_) (e.g. a mutation matrix) with

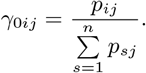

Hence, if a sample is perturbed in two genes, their responsibility for that sample is 50% each. However, another possibility is to include prior belief for the perturbations in a sample and not treat them equally. E.g. if perturbation *i* in sample *j* is twice as certain as perturbation *k*, we can set 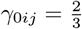 and 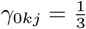.

## 3 Simulation study

As a proof of principle, we use NEM*π* to simulate data based on a ground truth. We simulate a data set based on random parameters: a causal network of the P-genes *φ*, E-gene attachments *θ* and a discrete perturbation matrix *Z*, which includes only the direct perturbations. The complete perturbation profile is computed by the network propagation Ω = *φ^T^ Z*. The simulations are based on *n* = 5, 10, 15 P-genes, 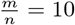 E-genes per P-gene and around 10 × *n* × 2 samples to ensure a reasonable amount of samples with correct perturbation profiles. In general, a higher number of E-genes decreases noise, especially because NEM*π* can exclude badly fitting E-genes. Additionally, we add 10% uninformative samples and E-genes which consist only of noise and are not related to the ground truth. As described in the Method section, the implementation of a null sample during iteration accounts for those samples. We allow roughly 20% double and 10% triple perturbations (columns in *Z* with more than one entry equal to 1). We add Gaussian noise with standard deviation *σ* = 1, 3, 5 for 100 runs each. We compare NEM*π* to support vector machines (svm, R-package e1071, Meyer *et al.*, 2019) neural networks (nn, Venables and Ripley, 2002), random forest (rf, Liaw and Wiener, 2002) and *k* nearest neighbours (Venables and Ripley, 2002) classification methods. We trained the classifiers on the labeled samples and computed the class label probabilities on the test and training set. We classify each sample and P-gene separately, i.e., we learn a classifier for each single P-gene based on the gene expression predicting whether the gene is perturbed in the sample or not. Afterwards we combine the class probabilities for each sample and P-gene to a matrix corresponding to the estimator of the perturbation profile 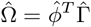 provided by NEM*π*. Additionally, we compare our results to the two data imputation methods, namely mice (Shah, 2018; Azur *et al.*, 2011) and missForest (Stekhoven and Bühlmann, 2011). We used the default implementations except for mice, which was running too long to converge. Hence, we reduced the number of iterations from 5 to 2.

We measured the degree of success as the area under the precision-recall curve (AUC, supplement) by comparing the ground truth perturbation profile Ω = *φ^T^ Z* with predicted perturbation profile 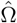.

### Samples with unobserved perturbation profiles

In a first study, we randomly removed the perturbation profiles for 10%, 50% and 90% of the samples and tried to infer them (Fig. 3). The AUC shows that we can recover the perturbation profiles very well, even at high noise levels and much better than the other methods. For 5 P-genes, random forest is the only competitive method. However, with only 10% unobserved samples, all methods do only marginally better than randomly guessing the unknown perturbations. With larger sets of unobserved samples, the classifiers stay mostly robust while random guessing drops in performance. For example, with 50% unobserved samples and medium noise for 10 P-genes, the best classifiers have an AUC of around 0.6, more than 0.1 above random guessing, while NEM*π* achieves an AUC of approximately 0.9. Additionally to the AUC, we also show the accuracy of the inferred network *φ*, (supplement, Fig. S1) and the E-gene attachment *θ* (supplement, Fig. S2). The accuracy of the network and attachment break down considerably at high noise levels, e.g., 50% network accuracy for 90% unobserved and *σ* = 5.

**Figure 3:**
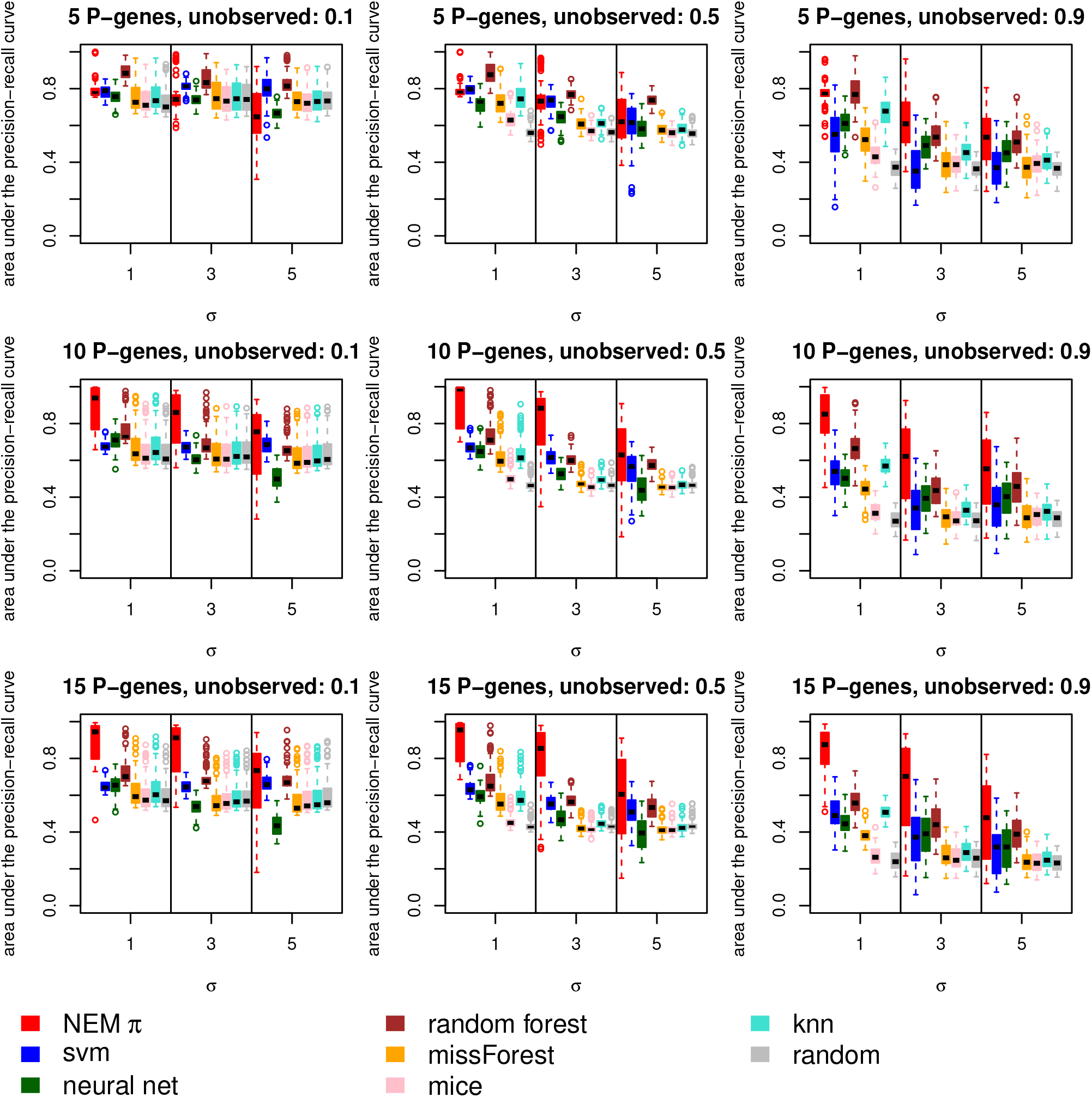
Unobserved perturbations. Shown is the area under the precision-recall curve between the predicted and ground truth perturbation profile. The columns show different amounts of samples with unobserved perturbation profiles (10%, 50%, 90%). The rows show different numbers of P-genes (5, 10, 15). Overall our approach (red) performs better than svm, neural nets, random forest, missForest, mice and k-nearest neighbours.

NEM*π* infers the network *φ* during optimization. However, if the underlying network is known, it can be provided and the network learning step is skipped. NEM*π* proves to be robust against false positive edges in the given network (supplement, Fig. S3), but less robust against false negatives (supplement, Fig. S4). We suggest to let NEM*π* do the network optimization unless there is high confidence in the prior network.

### Augmenting perturbation profiles

In a second study, we fixed the samples with unobserved perturbation profiles to 50% and randomly changed 10% and 50% of the rest of the perturbation profiles, respectively. For a random sample we first sampled the number of directly perturbed P-genes *x* ∈ {1, 2, 3} and then sampled *x* different P-genes to be perturbed. E.g. for a known sample we forget which genes are actually perturbed and draw *x* = 2. Hence, we sample two random P-genes and label the sample as perturbed by them instead. This study shows that we can successfully recover unobserved perturbations profiles, even when a fraction of the given perturbation profiles are incorrect (Fig. 4). Furthermore, we can even partially augment incorrect perturbations. For 15 P-genes and medium noise levels, NEM*π* has an AUC of over 0.8 when 75% are unobserved or incorrect.

**Figure 4:**
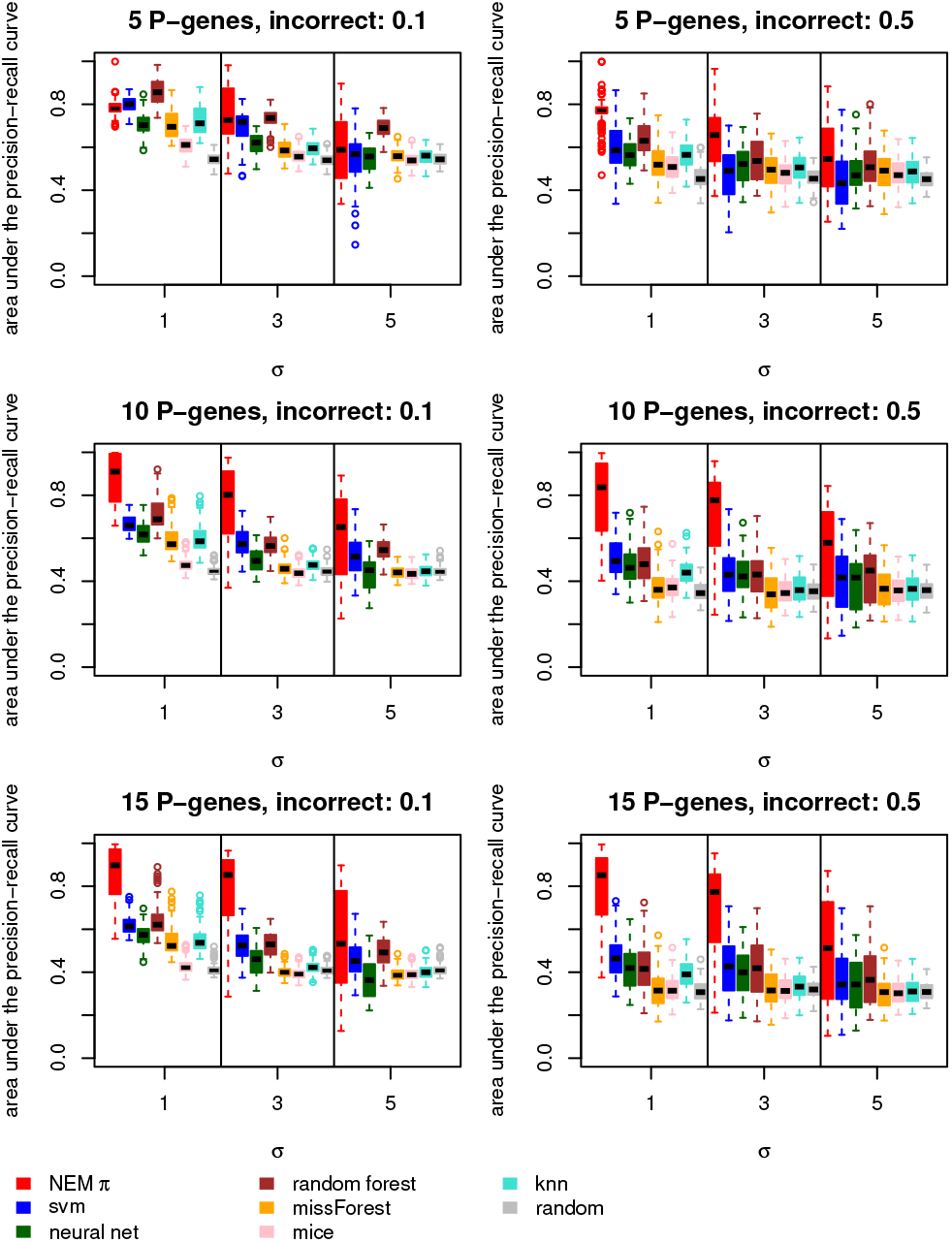
Unobserved and incorrect perturbations. We set the number of perturbation profiles unobserved at a constant 50%. Shown is the area under the precision-recall curve between the predicted and ground truth perturbation profile. The columns show different amounts of incorrect profiles (10%, 50%). The rows show different numbers of P-genes (5, 10, 15). Overall our approach (red) performs better than svm, neural nets, random forest, missForest, mice and k-nearest neighbours.

### Inference with respect to unknown P-genes

Lastly, we simulated data for a large number of P-genes *x* ∈ {50, 100}, but kept the number of informative samples fixed to 1000. However, we only included eight P-genes in our model and did not use any information about the other unknown P-genes during the inference. We wanted to investigate, how well we can infer the perturbation profiles of these eight P-genes. This is the most realistic scenario according to the results of Bailey *et al.*, 2018, who find eight driver genes exclusive to breast cancer and 64 pan-cancer driver genes. Together with non-pan-cancer non-exclusive genes, this makes a total of 93 perturbed genes.

The accuracy of the perturbation profiles decreases slightly due to the systematic noise of the unknown P-genes (Fig. 5). Interestingly, the accuracy is also more robust against Gaussian noise, due to the large sample size for only eight P-genes. The other methods break down completely. As would be expected, reduced sample sizes are diminishing to NEM*π*’s accuracy (supplement, Fig. S5-S6).

**Figure 5:**
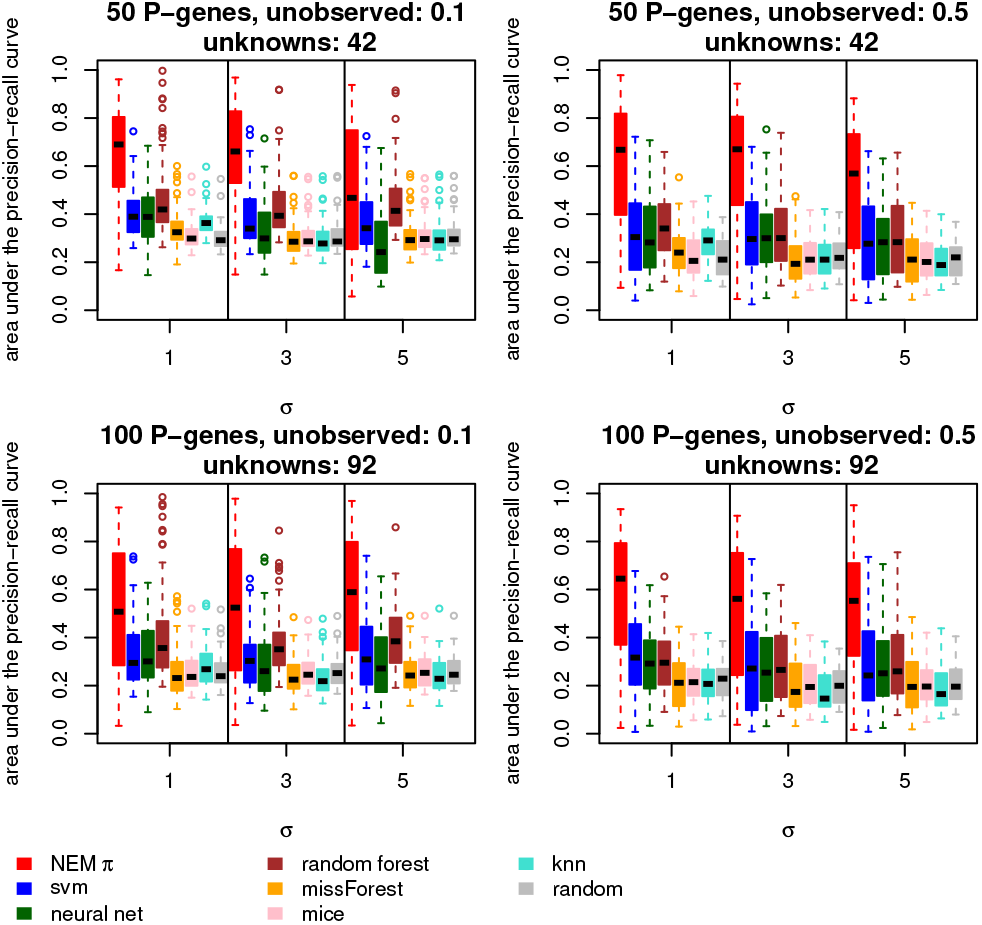
Unknown confounding P-genes. The area under the precision-recall curve between the predicted and ground truth perturbation profiles is shown in for 50 P-genes and 100 P-genes (rows), and 10% and 90% unobserved samples (columns), respectively. The number of known P-genes is set to eight.

## 4 Validation on CRISPR scRNA-seq data

We validate our approach on several CRISPR scRNA-seq data sets published by Adamson *et al.*, 2016 (GEO: GSE90546, Edgar *et al.*, 2002; Barrett *et al.*, 2012). Our goal is to predict the perturbations in all cells from a random subset. In this case, the perturbations have been introduced experimentally using Perturb-seq. The data sets are from a pilot study on 7 genes, a larger study on 82 genes, and an epistasis study on three genes. The last study consists of three data sets with different chemical treatments and includes double and triple perturbations. All genes are involved in the regulation of the endoplasmic reticulum pathway.

We removed genes with a median expression of zero counts and used the R package `Linnorm` (Yip *et al.*, 2017) to pre-process the single-cell data. For the computation of the log-odds, we refer to the supplement (p.1). After pre-processing, the five data sets consist of 1754 × 3927 (pilot), 2794 × 53290 (main), 3399 × 4015 (epistasis 1), 3615 × 3602 (epistasis 2) and 3614 × 3363 (epistasis 3) genes times cells.

After learning a network *φ* for each data set with the original NEM, we use the network as the ground truth for this validation study. We employ an exhaustive search for the data sets with three genes and greedy search for the others. The ground truth perturbation profile is computed by Ω = *φ^T^* Γ with Γ derived from the known cell labels, i.e., which P-gene has been perturbed in which cell.

For the validation, we remove the labels for 50% of the cells and use the different methods to re-learn the perturbations. For the main study, we randomly sample a subset of 10 and 15 P-genes. NEM*π* is not provided with the ground truth *φ* but has to learn the network from the partially labelled data (supplement, Fig S7). The accuracy is computed as before over 100 independent runs. All methods achieve the highest accuracy for the data sets from the epistasis studies (Fig. 6, bottom). For the other two data sets with more P-genes (7, 10, 15) the accuracy drops (Fig. 6, top). The main study shows the highest variation in accuracy due to the random sampling of P-genes. The accuracy is in general higher for 10% and lower for 90% unlabelled cells (supplement, Fig S8). We distinguished runs with a dense network inferred by NEM*π* and a sparse network (supplement, Fig S9). As expected, NEM*π* has more power for dense networks. Overall, NEM*π* achieves the highest accuracy and has the most success in predicting perturbations. The imputation method mice took too long to compute and did not converge.

**Figure 6:**
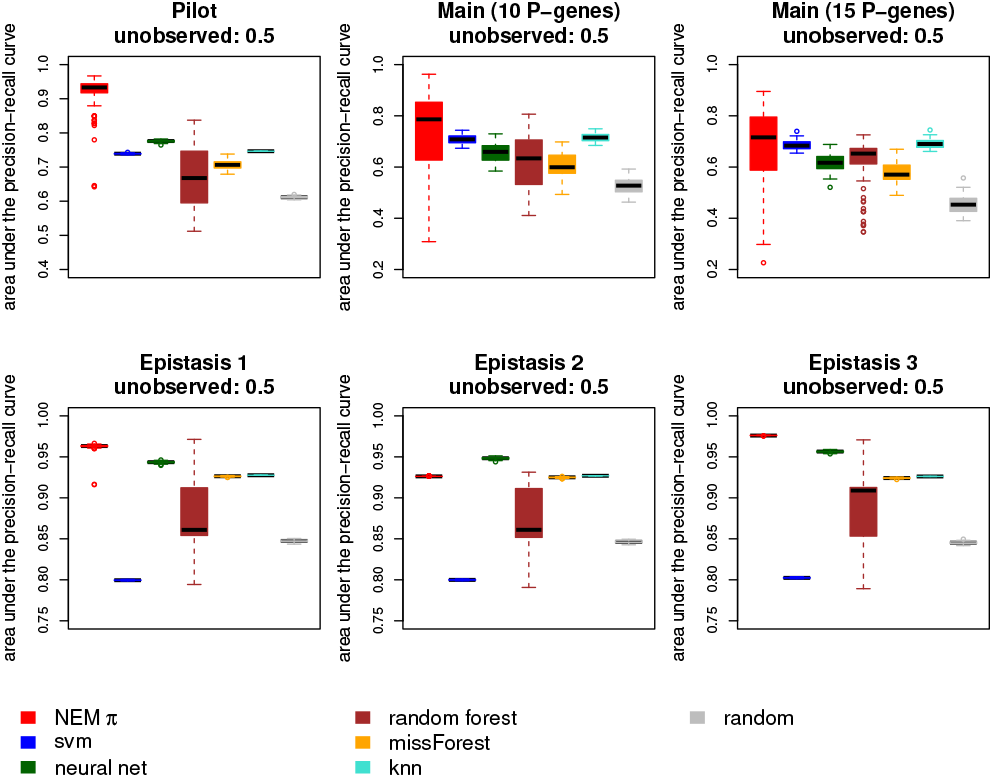
Accuracy of the various methods for the CRISPR scRNA-seq data sets. All methods perform very well for the epistasis studies except for svm (bottom). Overall NEM*π* outperforms the other methods, but shows a large variance over the randomly samples P-genes of the main study.

## 5 Exploratory analysis on breast cancer

We apply our method to the breast cancer (BRCA) data set from TCGA, which has many samples including controls. In this analysis, we want to explore the possibility to predict other perturbations, like copy number abberations or methylation. As the initial incomplete perturbation matrix, we use the mutation matrix *M* = (*m*_*ij*_) with *m*_*ij*_ = 1, if sample *j* has a mutation in gene *i* and otherwise *m*_*ij*_ = 0. We aim to 1) infer perturbation profiles for samples, which have no mutation data available (unobserved perturbations) and 2) augment the known mutations. As P-genes we choose the following driver genes previously identified as exclusive to BRCA (Bailey *et al.*, 2018): *CBFB*, *CDKN1B*, *GATA3*, *GPS2*, *MAP2K4*, *NCOR1*, *PTPRD*, *TBX3*.

We used the R-package `TCGAbiolinks` (Colaprico *et al.*, 2015) to access and download the gene expression counts and mutation information. We define a gene in a sample as mutated, if it was called by at least three of the four methods available in the TCGA data set (Fan *et al.*, 2016; Cibulskis *et al.*, 2013; Harris *et al.*, 2011; Koboldt *et al.*, 2012) to avoid false positives. Furthermore, we set mutations to 0, which were labeled as “silent= by TCGA.

The BRCA data set consists of 1215 samples including 113 control samples. We summarize duplicate samples and duplicate genes with the median. Roughly 92% of all samples do not carry a mutation for any of the P-genes, because either none were called or mutation data was not available. We filtered out lowly expressed genes (median < 10 counts) to obtain 20, 213 E-genes. We used edgeR (Robinson *et al.*, 2010) to normalise the gene expression. For the computation of the log odds we refer to the supplement. In the last step we removed uninformative E-genes (median log odds equal to 0) and were left with 19, 381 genes. The log odds for the BRCA data set follow a similar distribution than our simulated data. (supplement, Fig. S10).

NEM*π* takes roughly 2.5 minutes on a MacBook pro (2017) to converge in 42 iterations (supplement, Fig. S11). The inferred expectations Γ of the perturbation matrix *Z* is a sparse matrix with virtually binary predictions (Fig. 7, supplement, Fig. S11), i.e., many samples have only predicted 0s (white) for all except one P-gene (1, dark blue), which corresponds to a single direct perturbation. In only few samples the expectations show more uncertainty (light blue). All perturbations in Γ are propagated via the inferred causal network *φ* (Fig. 8). Hence, all samples with a direct perturbation of *MAP2K4* have also perturbations in all other P-genes except for *GATA3* and *TBX3*. Vice versa for all samples with a direct perturbation in *CDKN1B* our model predicts no perturbation in any other P-gene. Although, these eight genes have not been found to be in a joint signaling pathway, gene ontology analysis (Szklarczyk *et al.*, 2018, https://string-db.org/) shows evidence of similar activity in biological processes like the regulation of the B-cell receptor signaling pathway (supplement, Table S1).

**Figure 7:**
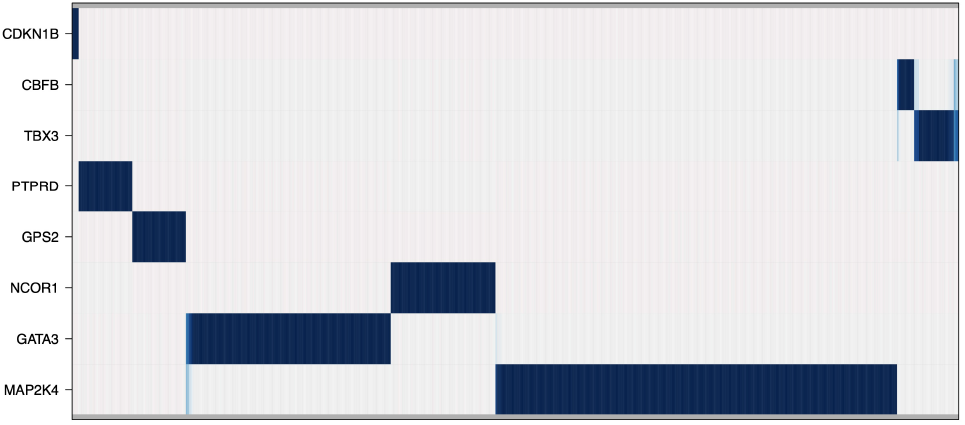
The expectations Γ of the direct perturbations *Z* inferred from the BRCA data set. Shown are the expectations of the P-genes (rows) for the samples (columns). Dark blue values are close to 1, while light blue values are between 0 and 1.

**Figure 8:**
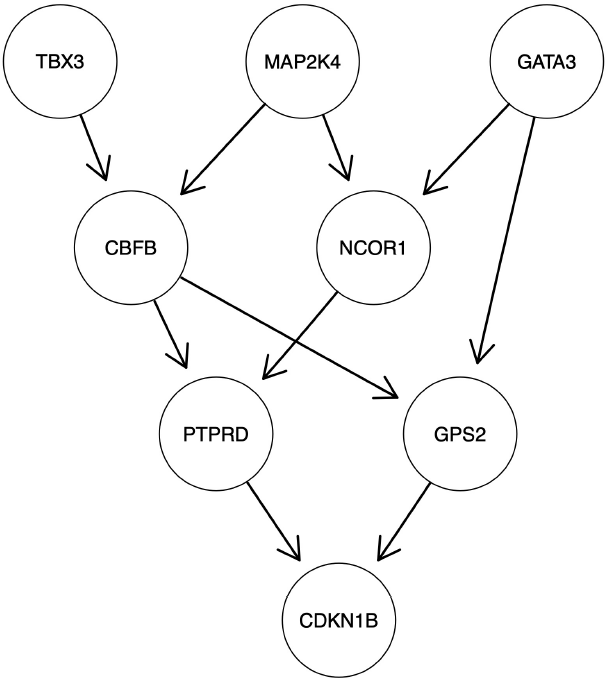
Causal network *φ* inferred from the BRCA data set. Shown is the causal network connecting the P-genes based on their effect on the gene expression. This network propagates perturbations predicted by Γ (Fig. 7) to all descendants. E.g. in all the samples with a perturbation of *MAP2K4* predicted by Γ all other genes except for GATA3 and *TBX3* are also perturbed.

### Comparison to other modes of perturbation

A perturbation of a gene can happen in different ways. For example, a gene is mutated in some samples and therefore perturbed on the DNA level. However, in other samples no mutation is observed, but a copy number aberration, which can also lead to a perturbation of the gene. Our predicted perturbation profiles for all samples (Ω) for the BRCA samples are learned based solely on observed mutations. To investigate whether we can capture other modes of perturbations, we compare our prediction to available copy number variation (CNV) and methylation data. CNVs are provided by TCGA as a matrix *C* with *c*_*ij*_ = 0, if there is no copy number aberration of gene *i* in sample *j*, and *c*_*ij*_ ∈ {−2, −1, 1, 2}, if there is a loss (−) or a gain, respectively. We binarized *C* by setting all entries ≠ 0 to 1. We called sites methylated (1) with a cutoff of > 0.5 for the beta score ∈ {0, 1} provided by TCGA. Those perturbations by methylations are stored in a matrix *H* with *h*_*ij*_ = 1, if gene *i* is methylated in sample *j*.

For visualisation, we binarized our predicted perturbation profile Ω with a cutoff of 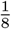 and combined the mutations matrix *M*, the CNV matrix *C*, the methylation matrix *H* and our predicted perturbations Ω in one matrix (Fig. 9). The dark blue regions show that our predictions based on the mutations capture unobserved perturbations implied by CNV or methylation not covered by mutations.

**Figure 9:**
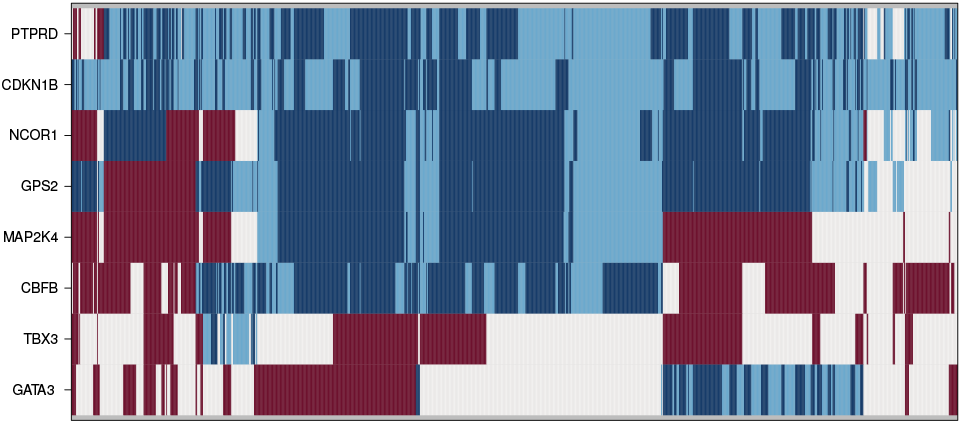
Predicted and measured perturbations. This matrix visualises predicted and measured perturbations (mutation, CNV or methylation) of the P-genes (rows) over all samples (columns). Dark blue are perturbations, which are predicted and confirmed by a measurement (true positives). Light blue perturbations are only predicted (false positives), red are only measured (false negatives) and white are neither (true negatives).

The high false positives (light blue) may be explained by the fact that we have not covered all types of perturbation. A gene can be indirectly perturbed without any CNV, mutation or methylation (e.g., micro RNA activity, Shivdasani, 2006; O’Brien *et al.*, 2018), or the aberration is not detected due to noise.

Even though *CDKN1B* is hardly mutated in any sample, we predict it as perturbed in all samples. It is downstream of all other genes in the network and hence perturbed, if any other gene is perturbed. *PTPRD* and *CDKN1B* have similar profiles with only few samples without a predicted perturbation.

*GATA3*, *MAP2K4* and *TBX3* on the other hand have the least amount of predicted perturbations. Additionally the predicted perturbations for all three genes are mutually exclusive. This is also reflected in the continuous matrix Γ (Fig. 7), where *GATA3*, *MAP2K4* and *TBX3* hardly share any samples (light blue).

Next, we used svm, random forest, neural net and k-nearest neighbours classifiers, and missForest to impute perturbations only from known mutations. For a comparison of the methods we computed the AUC of the precision-recall curve (Table 1, first row). The overall accuracy, while greater than random, is low. However, if we assume that the perturbations caused by CNVs and methylation are propagated by the network *φ* inferred by NEM*π*, the accuracy increase, especially for NEM*π* and svm (Table 1, second row).

**Table 1:** Area under the precision-recall curve for the eight breast cancer-specific driver genes. All methods achieve a marginally higher accuracy than random guessing in predicting CNVs and/or methylated sites (first row). If perturbations by CNVs and methylations are propagated by the network *φ* inferred by NEM*π*, the accuracy increases, especially for NEM*π* and svm (second row).

| NEMπ | svm | neural net | random forest | missForest | knn | random |
| --- | --- | --- | --- | --- | --- | --- |
| 0.60 | 0.62 | 0.58 | 0.63 | 0.60 | 0.59 | 0.54 |
| 0.88 | 0.87 | 0.76 | 0.78 | 0.74 | 0.78 | 0.74 |

Next, we randomly sampled 10 genes from the pan-cancer list of Bailey *et al.*, 2018 and predicted CNVs and methylation from mutations and gene expression profiles to asses the variance of the AUC over the data set. Overall the performance stays low (supplement, Fig. S12, left) for all methods with NEM*π* having a significantly greater accuracy than the other methods except for random guessing, which is still worse than NEM*π* but misses the 5% cut for significance (rank sum test of the accuracy values with alternative ‘greater’, p-value 0.06757). However, if the network *φ* inferred by NEM*π* is true, the perturbations caused by CNVs or methylations are propagated via the network *φ*. In this case, NEM*π* is also significantly better than random (supplement, Fig. S12, center). Additionally, NEM*π* is also almost 50 times faster than neural nets and random forest and 5 times faster than svm. Only missForest and knn are faster than NEM*π* (supplement, Fig. S12, right).

In an additional analysis, we performed leave-one-out cross-validation exclusively on mutated samples. We removed one sample and trained a model on the mutation and expression profiles of the remaining samples. Then we predicted the mutation profile of the removed sample solely based on its expression profile. NEM*π* achieves the highest AUC (supplement, Fig. S13). However, overall accuracy is very low across all methods. This may be explained by the fact that all methods try to predict perturbations, including, for example, CNVs and methylated sites, and not only mutations, which can be sparse.

While NEM*π* is not designed to predict driver genes, there might be an overlap between drivers and highly perturbed genes identified by NEM*π*. To asses a potential overlap, we compare the previous results (Fig. 9) with the driver gene identification method DawnRank (Hou and Ma, 2014). However, DawnRank does not build a predictor like the other methods. Instead it uses mutation calls, gene expression data, and a prior gene network to infer sample-specific driver gene rankings, normalized to a range of 0 to 1. Therefore, we apply DawnRank to the mutations, gene expression data, and a prior network from String-db (Szklarczyk *et al.*, 2018). We used the pre-processed network from the R package Prodigy (Dinstag and Shamir, 2019). We compare the normalized ranks of our genes of interest to the mutation calls just like in the leave-one-out cross-validation with the AUC of the precision-recall curve. DawnRank achieves an AUC of 0.15. However, DawnRank predicts *CDKN1B* as a driver gene with a score of close to 100% in all samples (minimum: 99.72%, Fig. S14). This agrees with the perturbation pattern of *CDKN1B* predicted by NEM*π* (Fig. 9). Hence, for the eight driver genes, *CDKN1B* is the most significant gene in both analyses and adds support to our assumption that driver genes and highly perturbed genes overlap.

## 6 Discussion

We have introduced NEM*π*, a novel method for inferring perturbation profiles of biological samples based on known incomplete perturbations and gene expression data. We have shown in a simulation study that our method successfully learns perturbation profiles in several different situations with the help of the underlying causal network of the perturbed genes. Overall, we achieve higher accuracy than comparable methods usually used for such a problem (i.e., support vector machines, neural networks, random forest, k-nearest neighbours, and the imputation methods missForest and mice). However, these methods are at a disadvantage, because they do not assume an underlying network, which is used to generate the data.

We applied NEM*π* to several CRISPR scRNA-seq data sets. This allowed for a validation of NEM*π* on real data in a controlled setting. We show that NEM*π* achieves high accuracy in learning the removed labels (known perturbations) from a subset of cells. Especially on the data set with only three genes and combinatorial perturbations, all methods achieve high accuracy. The accuracy decreases for the data sets with more perturbed genes. Naturally, on the data set with a random sample of 10 and 15 P-genes over the 100 runs the variance increases due to the heterogeneity of the sampled cells and the inferred underlying network.

We applied our method to the breast cancer (BRCA) data from TCGA. We chose the data set for its large number of samples, including controls. Control samples are necessary to normalize the tumor samples with respect to differential expression. With few or no control samples, normalization becomes more difficult and unreliable. We chose to infer the perturbation profiles of eight genes, which have been previously identified as exclusive to breast cancer. Hence, the genes should be highly relevant and our simulations have shown that we can account for unknown P-genes. Furthermore, we selected the P-genes in the application especially since it is known that they are driver genes, which are unique to BRCA. Hence, it is expected that those genes are highly perturbed in that cancer type.

We learned the perturbation profiles purely from mutations (and gene expression) data. We compared our predictions to other data sets implying gene perturbations (CNV, methylation). This shows that NEM*π* recovers many perturbations not included in the mutation profiles. However, there are also predictions with no measured perturbations and vice versa. The reason for this diversion can be simply noise. False negative predictions can also occur, because some genetic aberrations do not cause a perturbation of the gene. False positive predictions can be explained by an indirect perturbation of gene by other means than genetic aberrations (propagated perturbation). For randomly sampled P-genes (known pan-cancer genes), NEM*π* achieves on average the largest area under the precision-recall curve. If we assume that perturbations caused by CNVs and methylations are propagated by the network *φ* inferred by NEM*π*, the accuracy increases.

As shown in the application to simulated and real data, NEM*π* can handle a high number of samples and E-genes. Prediction accuracy is robust even against a large underlying network with only few known P-genes. While NEM*π* can be applied to more than 15 P-genes, accuracy decreases and run-time increases. Hence, we suggest to employ other methods to reduce the amount of P-genes to a feasible number. It would be interesting to extend NEM*π* with a divide and conquer approach to make it applicable to a larger set of genes. E.g. NEM*π* could be applied to subsets of genes to predict local perturbation profiles. How those profiles would be summarized still remains an open problem.

While we use the causal network to predict indirect perturbations, interpretations of the network itself is not clear. The network is build based on similar expression profiles among the P-genes. However, it is not clear, if the P-genes are actually directly or indirectly interacting with each other.

