## Supplement for "Inferring perturbation profiles of cancer samples"

1 Department of Biosystems Science and Engineering, ETH Zurich,
Basel, 4058, Swizerland and
2 Swiss Institute of Bioinformatics, Basel, 4058, Switzerland.
* To whom correspondence should be addressed.

December 10, 2020

**Log odds computation** We use the R package `ks` (Duong, 2019) to estimate the cumulative distribution function with a Gaussian kernel for control $f_c$ and tumor samples $f_t$ for each gene $i$. For each gene expression value $g_{ij}$ of a sample $j$, we computed

$$r_{ij} = \log \left( \frac{\min \{1 - f_t(g_{ij}), f_t(g_{ij})\}}{\min \{1 - f_c(g_{ij}), f_c(g_{ij})\}} \right).$$

We compared the distribution of the normalized TCGA breast cancer and simulated data (Fig. S8).

**Area under the precision-recall curve** We look at the area under the precision-recall curve for each simulation run. We choose 100 equidistant values between 0 and 1 as the respective cutoffs for the probabilities. For each cutoff, we compute precision and recall by

$$\text{precision} = \frac{tp}{tp + fp} \text{ and recall} = \frac{tp}{tp + fn} \tag{S1}$$

with $tp$, $fp$ and $fn$ as the amount of true/false positives and false negatives, respectively. These give the discrete curve with recall on the x- and precision on the y-axis. Of this curve we compute the area below over the 100 simulation runs.

**Accuracy of $\phi$ and $\theta$** We assess the accuracy of the network $\phi$ with a normalised hamming distance ham between the inferred and the ground truth network computed by

$$\mathcal{H} = 1 - \frac{\text{ham}(a, b)}{n(n-1)} \tag{S2}$$

with $n(n-1)$ as the maximum hamming distance for $n$ P-genes (Fig. S1). We assess the accuracy of $\theta$ by the total number of correctly attached E-genes. E.g., if 50 of 100 E-genes are correctly assigned, the accuracy is 50% (Fig. S2).

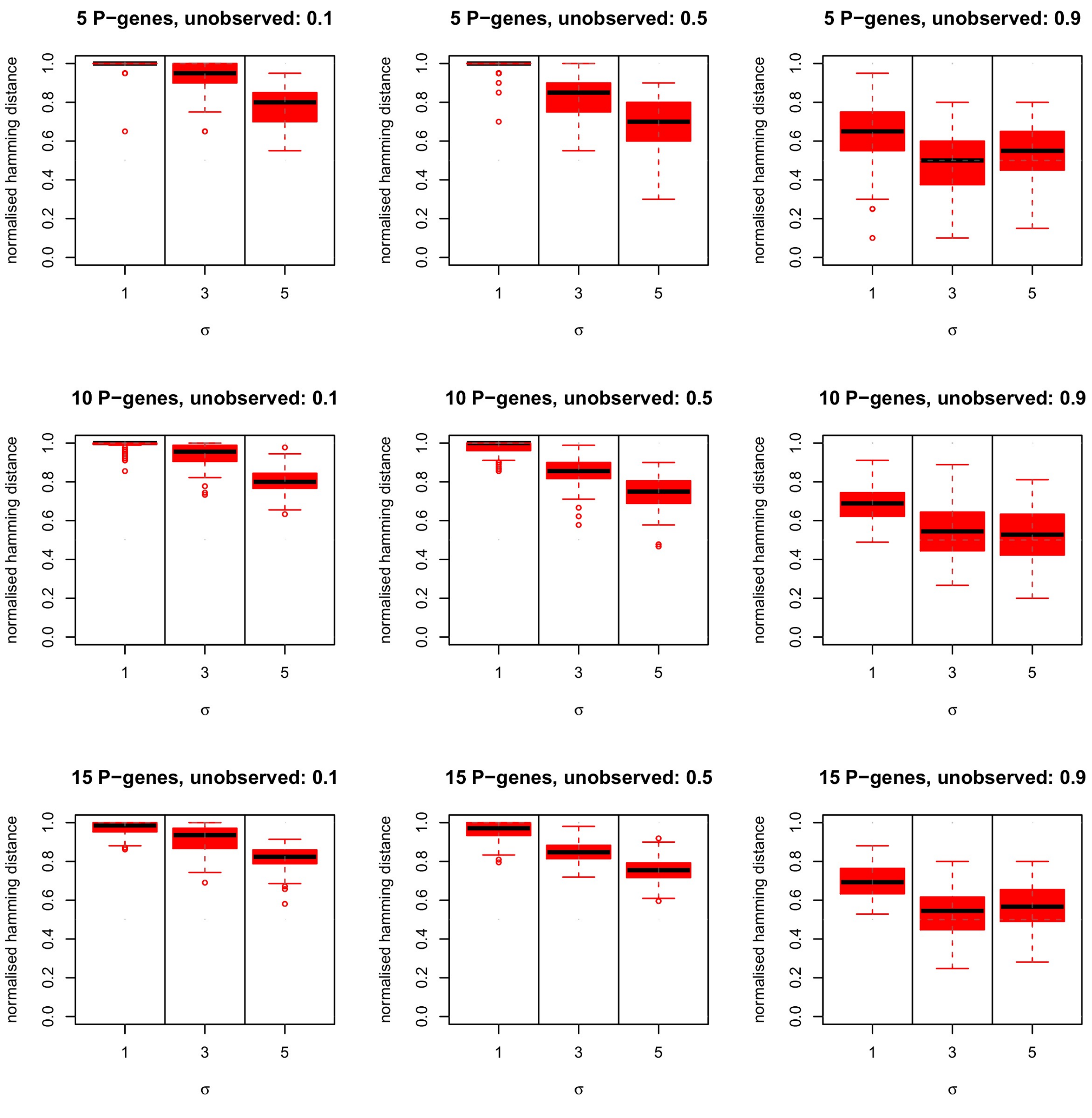


Figure S1: Network accuracy (Eq. S2) analogous to Fig. 3 in the main text.

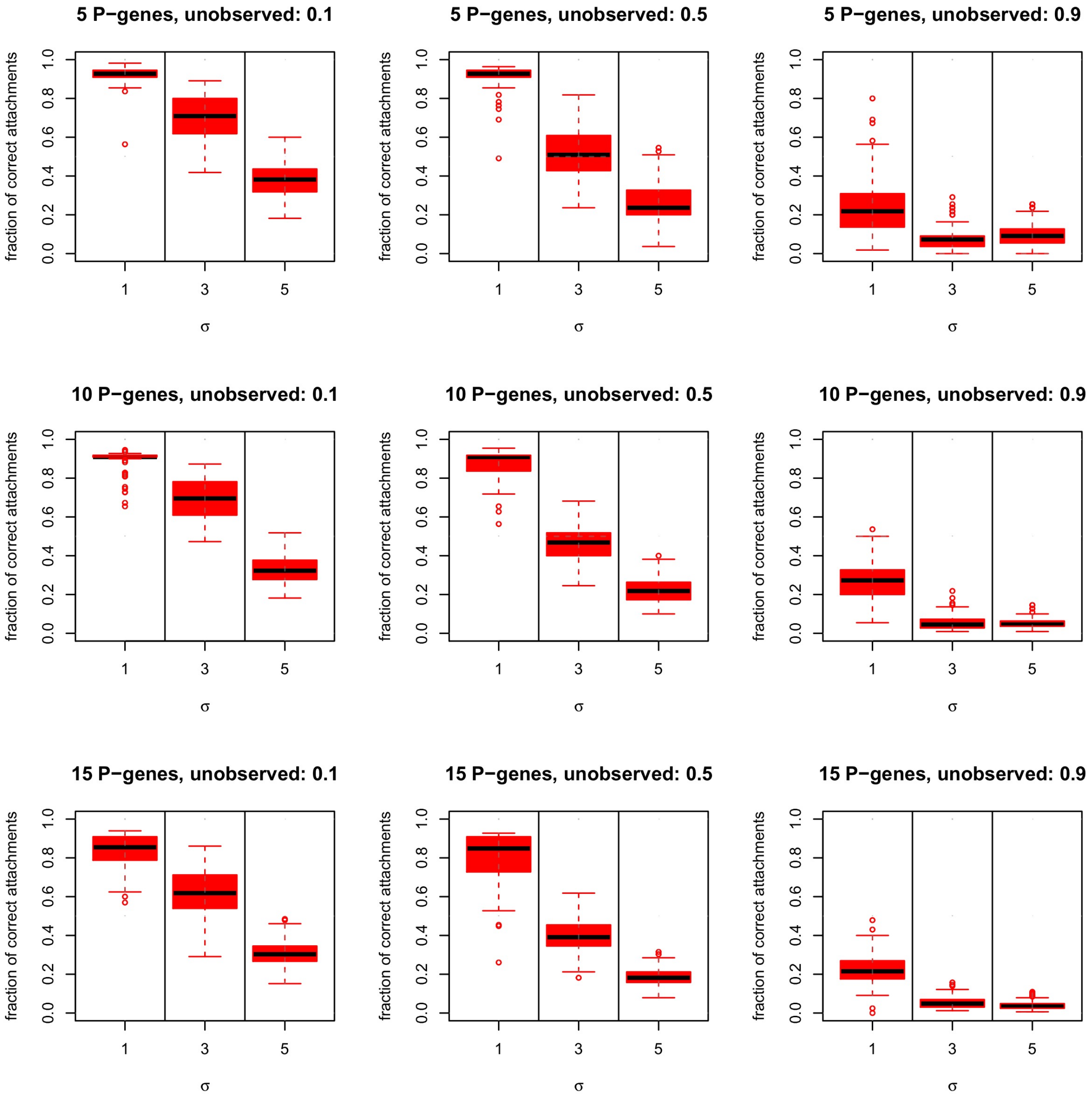


Figure S2: Accuracy of attachments $\theta$ analogous to Fig. 3 in the main text.

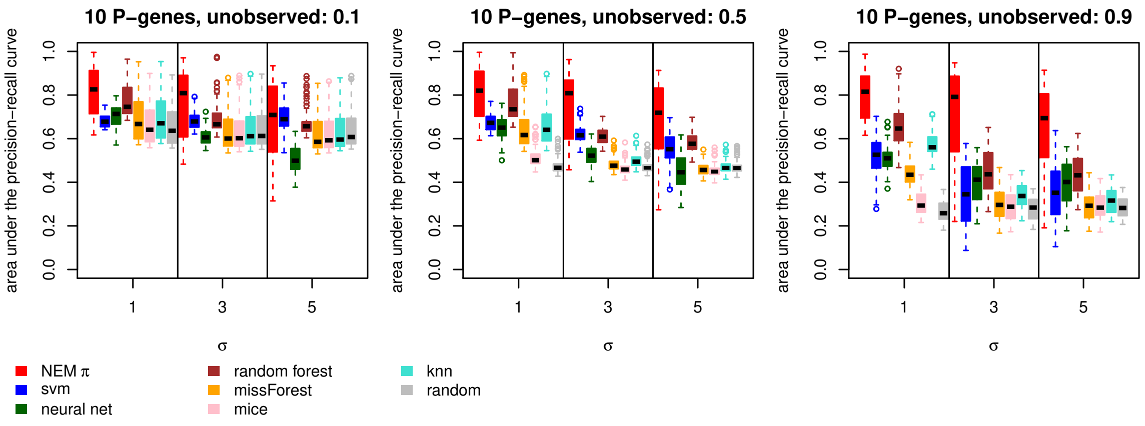


Figure S3: Accuracy of the perturbations profile analogous to Fig. 3 in the main text with 50% of additional incorrect edges.

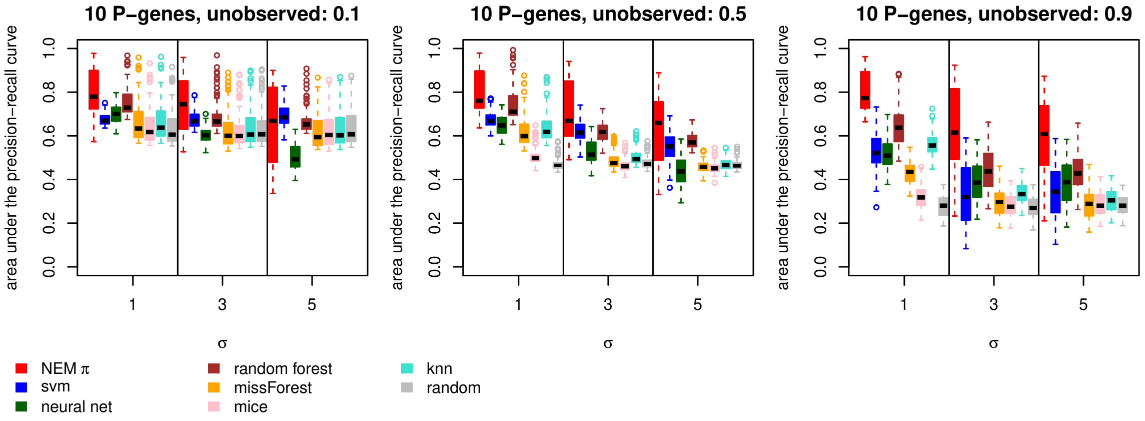


Figure S4: Accuracy of the perturbations profile analogous to Fig. 3 in the main text with 50% of edges removed.

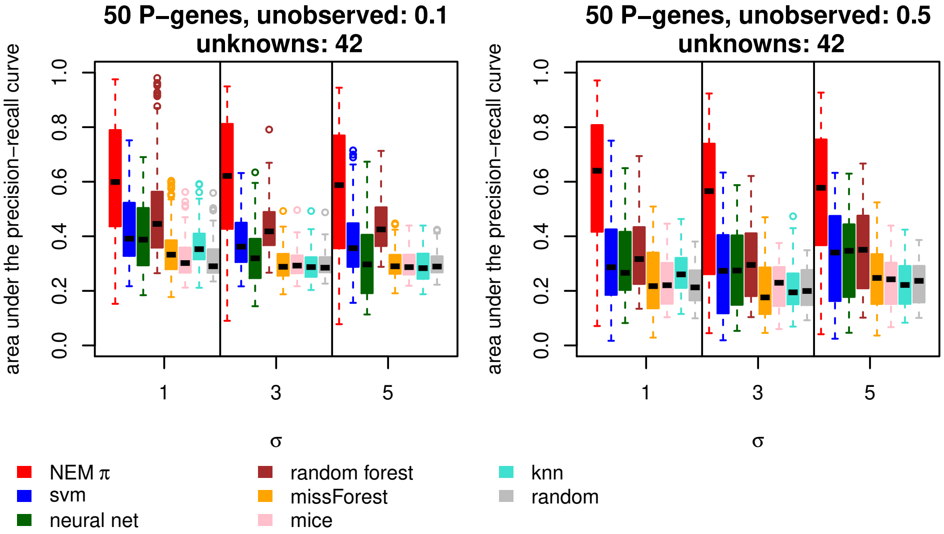


Figure S5: Accuracy of the perturbations profile analogous to Fig. 5 in the main text with 500 instead of 1000 samples.

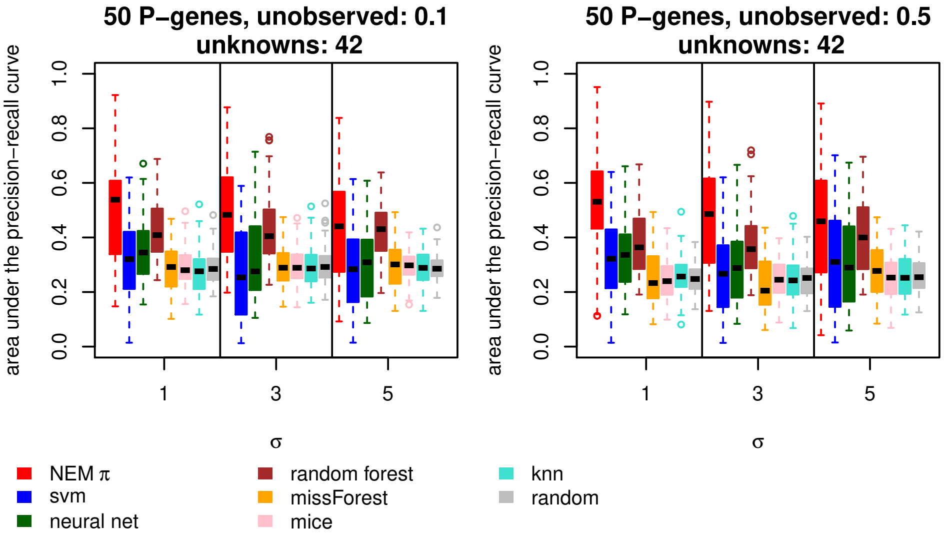


Figure S6: Accuracy of the perturbations profile analogous to Fig. 5 in the main text with 100 instead of 1000 samples.

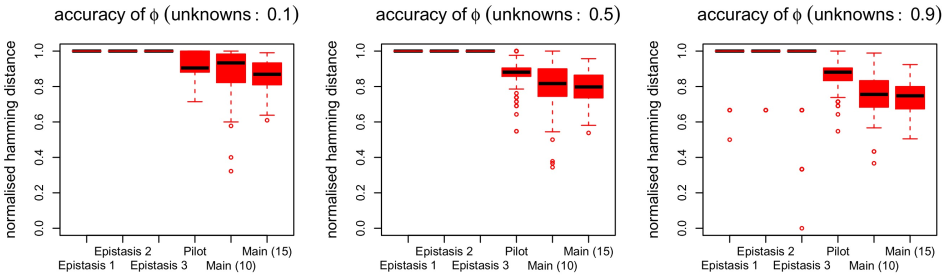


Figure S7: Network accuracy (Eq. S2) of $\phi$ inferred by NEM$\pi$ in the CRISPR validation study.

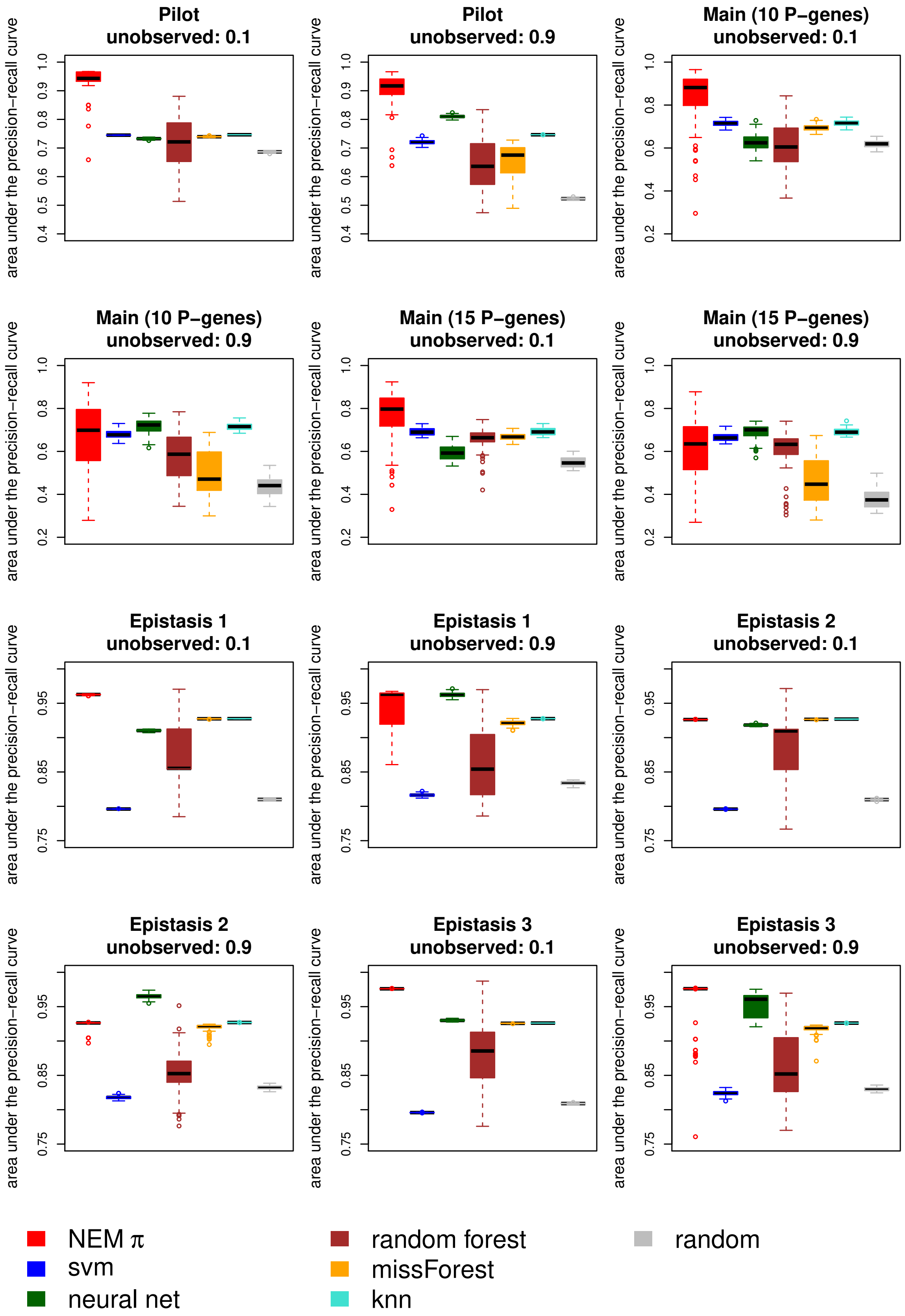


Figure S8: Accuracy of all methods for the CRISPR validation study with 0.1 and 0.9 unlabelled cells.

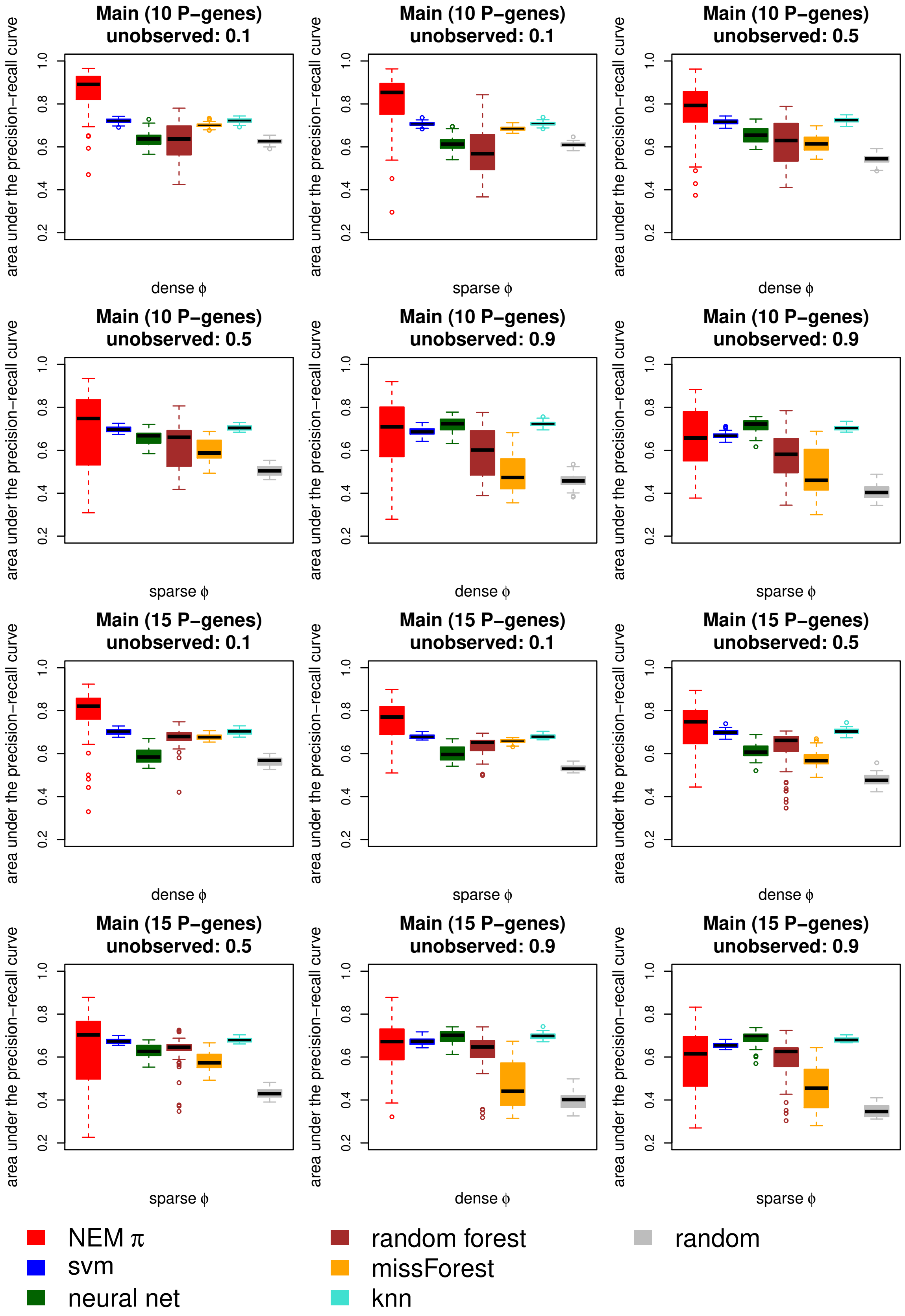


Figure S9: Accuracy of all methods for the CRISPR main validation study. The runs were split up into dens and sparse ground truth networks *phi*.

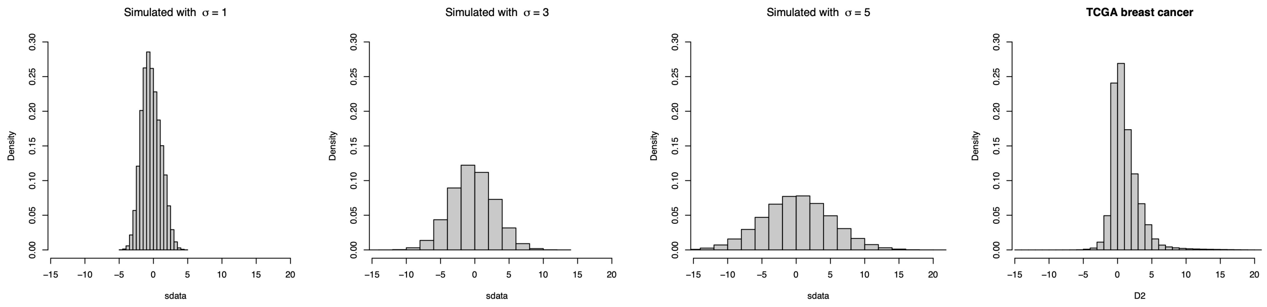


Figure S10: Simulated vs TCGA data: The three histograms on the left show the log odds distribution of simulated data for the three levels of noise ($\sigma = 1, 3, 5$). The right histogram shows the log odds distribution of the normalised data from TCGA. Our simulations show a similar or more extreme (noisy) distribution of log odds.

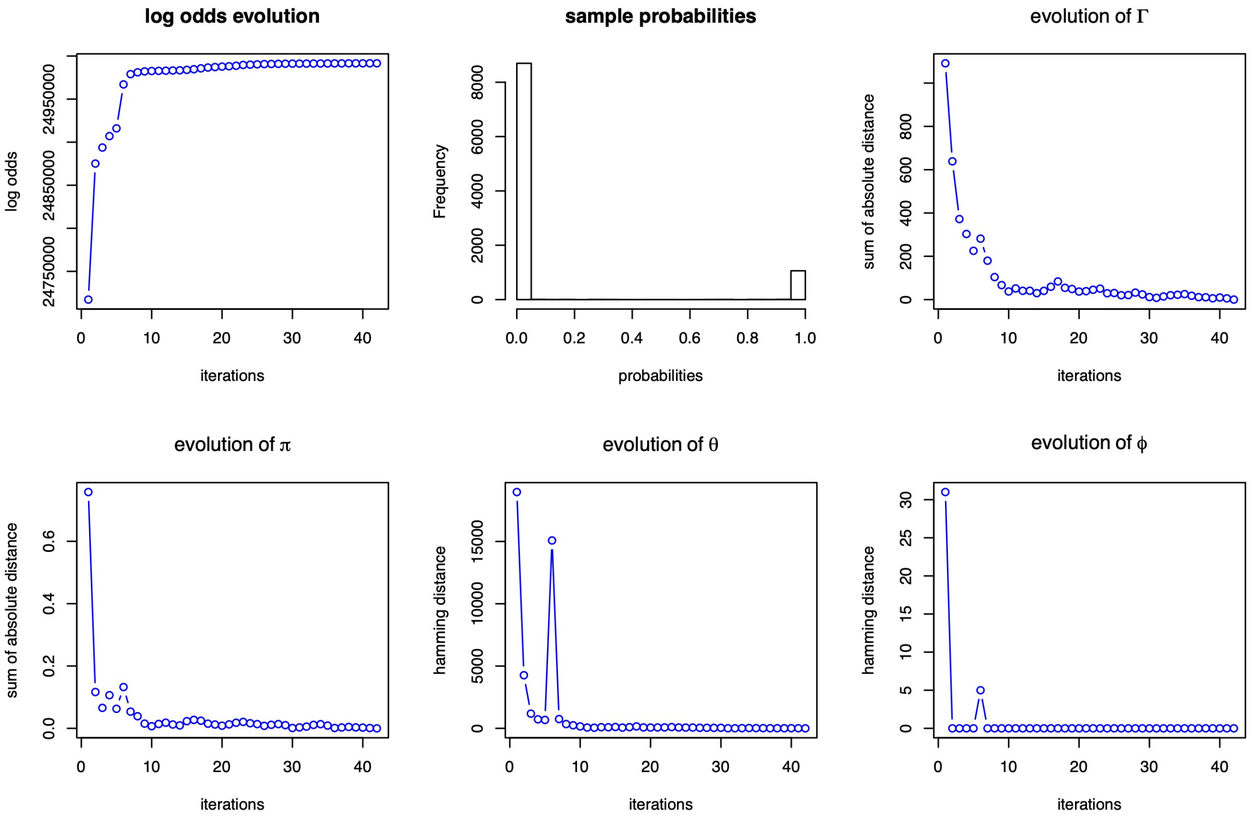


Figure S11: Convergence plots. Different plots showing the convergence of the NEM$\pi$ algorithm and the class probabilities (top, center). NEM$\pi$ took roughly 2.5 minutes to converge on a MacBook pro (2017). The convergence is shown by the log likelihood ratio (top, left), the perturbation matrix $\Gamma$ (top, right), the mixture weights $\pi$ (bottom, left), the E-gene attachments $\theta$ (bottom, center) and the P-gene network $\phi$ (bottom, right).

**Enrichment of biological processes** We used string-db (Szklarczyk *et al.*, 2018, https://string-db.org/) to look for enriched biological processes (Gene ontology, GO) connecting the genes in our network.

| Biological process | Genes | GO Gene set size | P-value |
|---|---|---|---|
| regulation of epithelial cell differentiation | CBFB, CDKN1B, GATA3, TBX3 | 137 | 0.0002 |
| regulation of B cell receptor signaling pathway | CBFB, GPS2 | 22 | 0.0023 |
| negative regulation of JNK cascade | GPS2, NCOR1 | 33 | 0.0038 |
| cardiac muscle cell development | MAP2K4, TBX3 | 49 | 0.0047 |

Table S1: Table of biological processes significantly enriched with a subset of the analysed P-genes.

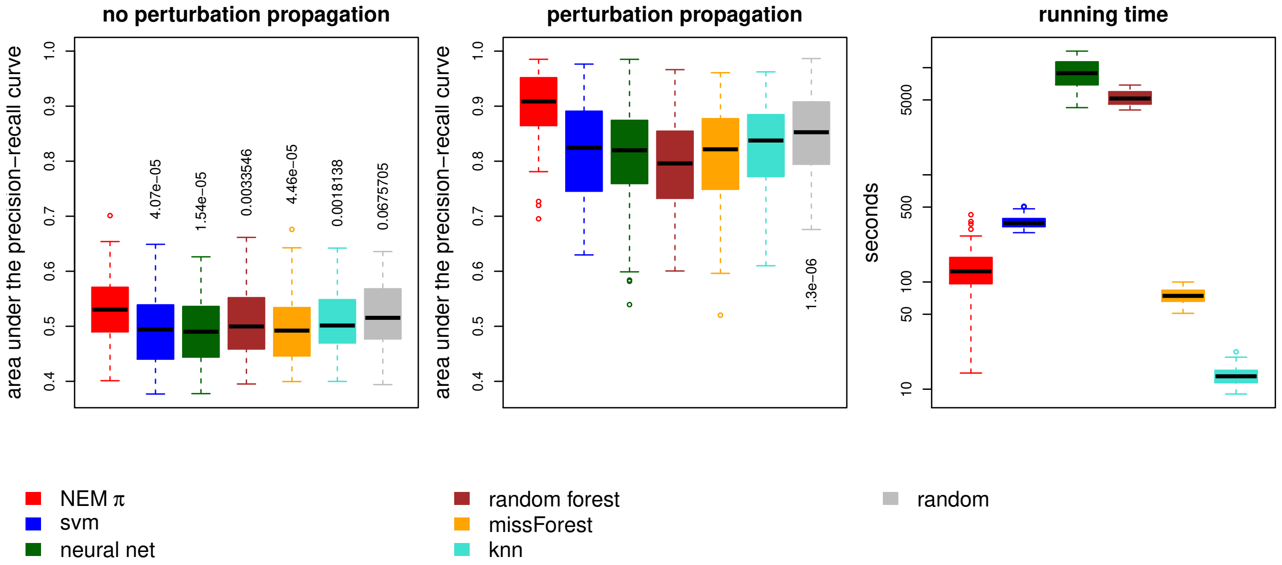


Figure S12: Accuracy for randomly sampled P-genes (pan-cancer drivers) from the TCGA breast cancer data. If we compare the predicted perturbations with CNVs and methylations (left), the area under the precision-recall curve is low for all methods, which are even lower than random guessing except for NEM$\pi$. If we compare the predictions to the perturbations propagated by the network $\phi$ inferred by NEM$\pi$ (center), the accuracy increases for all methods and NEM$\pi$ is significantly better than random. Additional NEM$\pi$ is faster than every method except for knn and missForest (right).

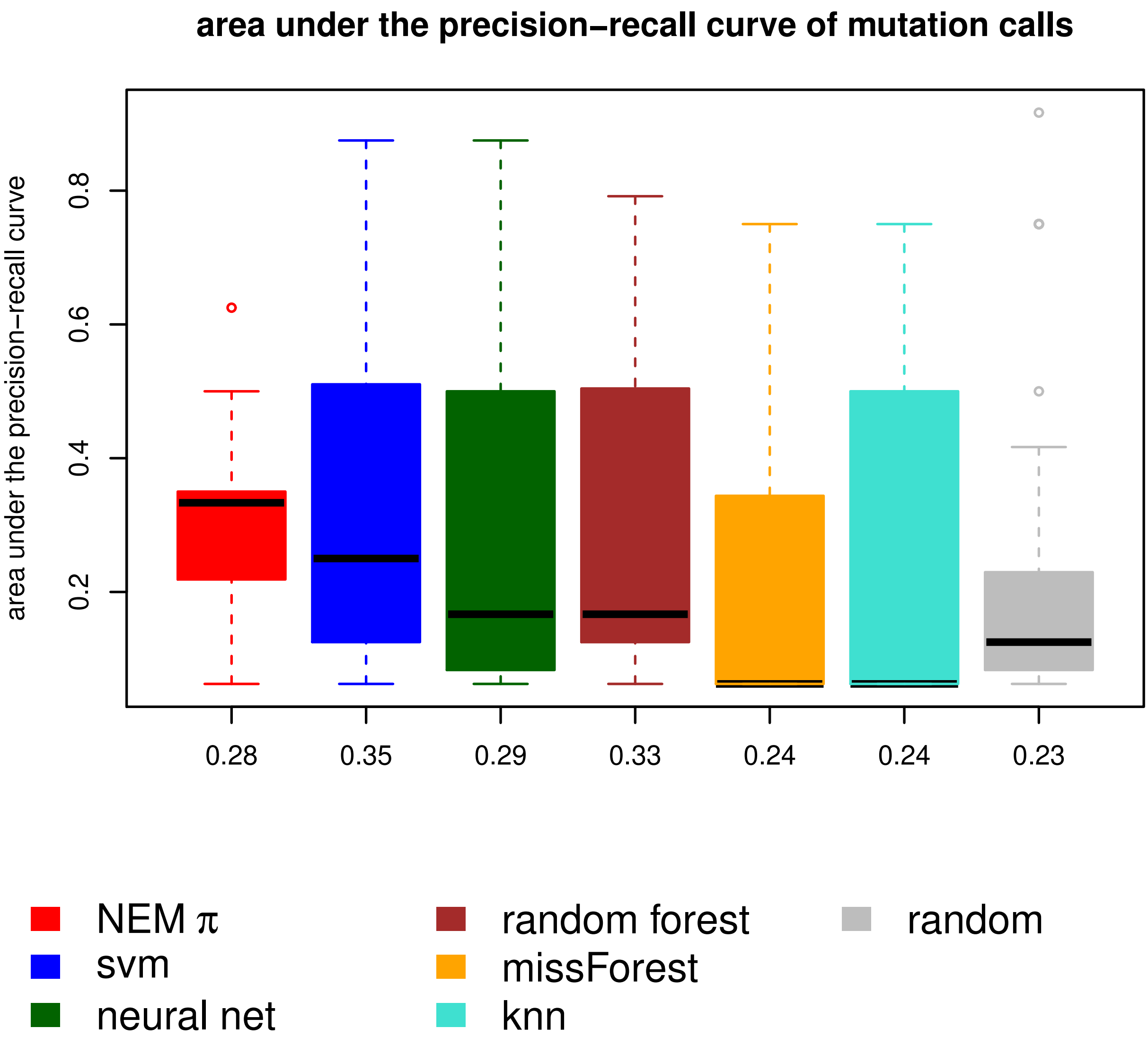


Figure S13: Boxplots of the area under the precision-recall curve for each leave one out cross validation over all 91 mutated samples. The means are shown on the x-axis. NEM$\pi$ (red) has the highest median. However, SVM (blue), neural nets (green) and random forest (brown) achieve a similar accuracy with higher variance.

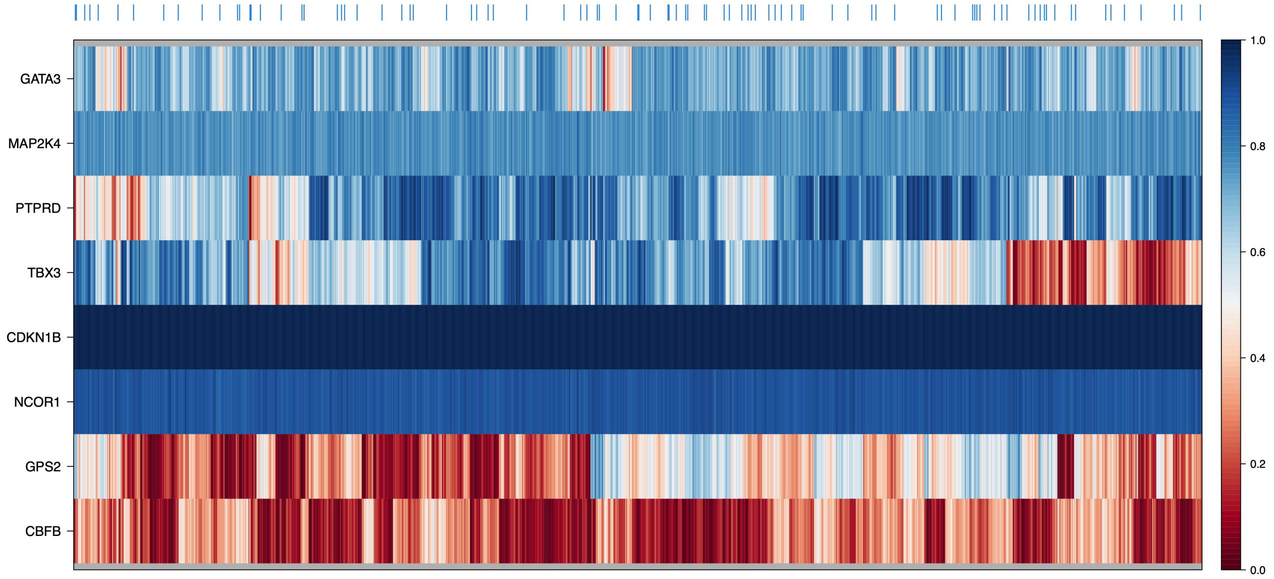


Figure S14: Heatmap of the normalized ranks from the driver gene prediction by DawnRank. Samples with mutations are marked with a blue tick above the heatmap. *CDKN1B* is the most significant P-gene over all samples.
